# The Nucleome of Developing Murine Rod Photoreceptors

**DOI:** 10.1101/369702

**Authors:** Issam Al Diri, Marc Valentine, Beisi Xu, Daniel Putnam, Lyra Griffiths, Marybeth Lupo, Jackie Norrie, Jiakun Zhang, Dianna Johnson, John Easton, Ying Shao, Victoria Honnell, Sharon Frase, Shondra Miller, Valerie Stewart, Xiang Chen, Michael A. Dyer

## Abstract

The nuclei of rod photoreceptors in mice and other nocturnal species have an unusual inverted chromatin structure: the heterochromatin is centrally located to help focus light and improve photosensitivity. To better understand this unique nuclear organization, we performed ultra-deep Hi-C analysis on murine retina at 3 stages of development and on purified rod photoreceptors. Predicted looping interactions from the Hi-C data were validated with fluorescence in situ hybridization (FISH). We discovered that a subset of retinal genes that are important for retinal development, cancer, and stress response are localized to the facultative heterochromatin domain. We also used machine learning to develop an algorithm based on our chromatin Hidden Markov Modeling (chromHMM) of retinal development to predict heterochromatin domains and study their dynamics during retinogenesis. FISH data for 264 genomic loci were used to train and validate the algorithm. The integrated data were then used to identify a developmental stage– and cell type-specific core regulatory circuit super-enhancer (CRC-SE) upstream of the *Vsx2* gene, which is required for bipolar neuron expression. Deletion of the *Vsx2* CRC-SE in mice led to the loss of bipolar neurons in the retina.

---

We have previously analyzed DNA methylation [whole-genome bisulfite sequencing (WGBS)], histone modifications [chromatin immunoprecipitation with DNA sequencing (ChlP-seq) for 8 marks], chromatin accessibility [assay for transposase-accessible chromatin using sequencing (ATAC-seq)], and transcriptional regulation through enhancer interactions (Brd4, CTCF, RNA-PolII, and ChIP-seq) in detail on the developing murine retina^1^. By integrating those data with RNA-seq data, we identified putative developmental stage– and cell type–specific enhancers, superenhancers, and CRC-SEs^1,2^. Many of those regulatory elements are located far away (10-200 kb) from their promoters and would require looping of the intervening DNA to regulate gene expression^1^.

Some studies have suggested that that the 3-dimensional (3D) chromatin landscape is relatively stable and that many long-range enhancer–promoter interactions are present, even before the gene is activated^3^. In contrast, others have found that gene activation by distal enhancers is accompanied by changes in the 3D chromatin landscape^4–7^. In addition to these putative dynamic looping interactions during retinal development, it was recently shown that some genes are sequestered in facultative heterochromatin in a cell type– and developmental stage–specific manner in the developing retina^1,8^. This is particularly notable in murine rod photoreceptors due to their unique nuclear organization.

Most nuclei have diffuse, transcriptionally active, and gene-rich euchromatin located in their centers, and more condensed heterochromatin is usually located on the nuclear lamina. Rod photoreceptors of nocturnal species have an inverted nuclear organization, with centrally located heterochromatin^9^. This pattern of chromatin optimizes vision in low light by reducing light scattering as it passes through the outer nuclear layer^9^. The difference in refractive index of euchromatin, relative to heterochromatin, combined with the inverted pattern creates a series of aligned converging lenses in the outer nuclear layer of nocturnal species. Therefore, in addition to the importance of nuclear organization in gene regulation, this is an example of the substantial influence of nuclear organization on cell and tissue adaptation and function.

Here we extend our previous analysis of chromatin accessibility and histone and DNA modifications during retinal development to include higher-order chromatin organization via ultra-deep Hi-C analysis of developing mouse retina and purified rod photoreceptors. We also developed an algorithm based on our integrated epigenetic and Hi-C analysis to predict localization to euchromatin and heterochromatin domains in retinal progenitor cells and rod photoreceptors. All data have been integrated into the following freely available cloud-based viewer: https://pecan.stjude.cloud/retinalnucleome. We used this integrated viewer to identify the first developmental stage– and cell type–specific CRC-SE that is required for the development of bipolar cells.

## Mapping chromatin interactions during retinogenesis

To elucidate cell- and developmental stage–specific promoter–enhancer interactions and other dynamic topologic features of the retinal genome, we performed Hi-C on embryonic day (E) 14.5, postnatal day (P) 0, adult murine retinae, and green fluorescent protein-positive (GFP^+^; rod photoreceptors) and GFP^-^ cells (cone, bipolar, horizontal and ganglion cells and Müller glia) from *Nrl-GFP* mice^10^. In total, more than 60 billion read pairs were sequenced across the 5 samples and compared to the 1.7 billion read pairs of Hi-C data published previously from the mouse cortex, fibroblasts, and murine embryonic stem cells (ESCs)^11^ (Table S1). The data were analyzed using Juicer^12^ and can be visualized on our viewer. We used multiple parameters and methods (Arrowhead^13^, Armatus^14^, and DomainCaller^11^) to identify topologically associated domains (TADs) and compare our data with previously published Hi-C data^11^ (Table S1). At the global level, the genome was dramatically reorganized during development. At E14.5, 2434 Mb of the genome were assigned to TADs, which was similar to that in adult mouse retina (2216 Mb). However, the number of TADs increased by 43% from 3690 at E14.5 to 5290 in adult retina, suggesting that domain boundaries were refined and subdivided with neuronal differentiation.

The deeper coverage of our dataset allowed us to identify regions in the retinal genome that were not mapped at high resolution in the previous analysis of mouse cortex and to identify contacts at 1-kb resolution (Table S1). For example, we identified a region on murine chromosome 4 with properties of a retinal-specific TAD, including several developmentally regulated genes and super-enhancers (Figs. 1A-C and S1). The region contains the *Icmt* gene encoding isoprenylcysteine carboxyl methyltransferase, which is required for photoreceptor function^15^, the *Kcnab2* gene encoding a potassium voltage-gated channel that is lost in 1p36-deletion syndrome^16,17^, and the *Rnf207* gene encoding a ring finger protein that is expressed only in 11-month-old mouse retinae (Figs. 1A and S1A,B).

**Figure 1.**
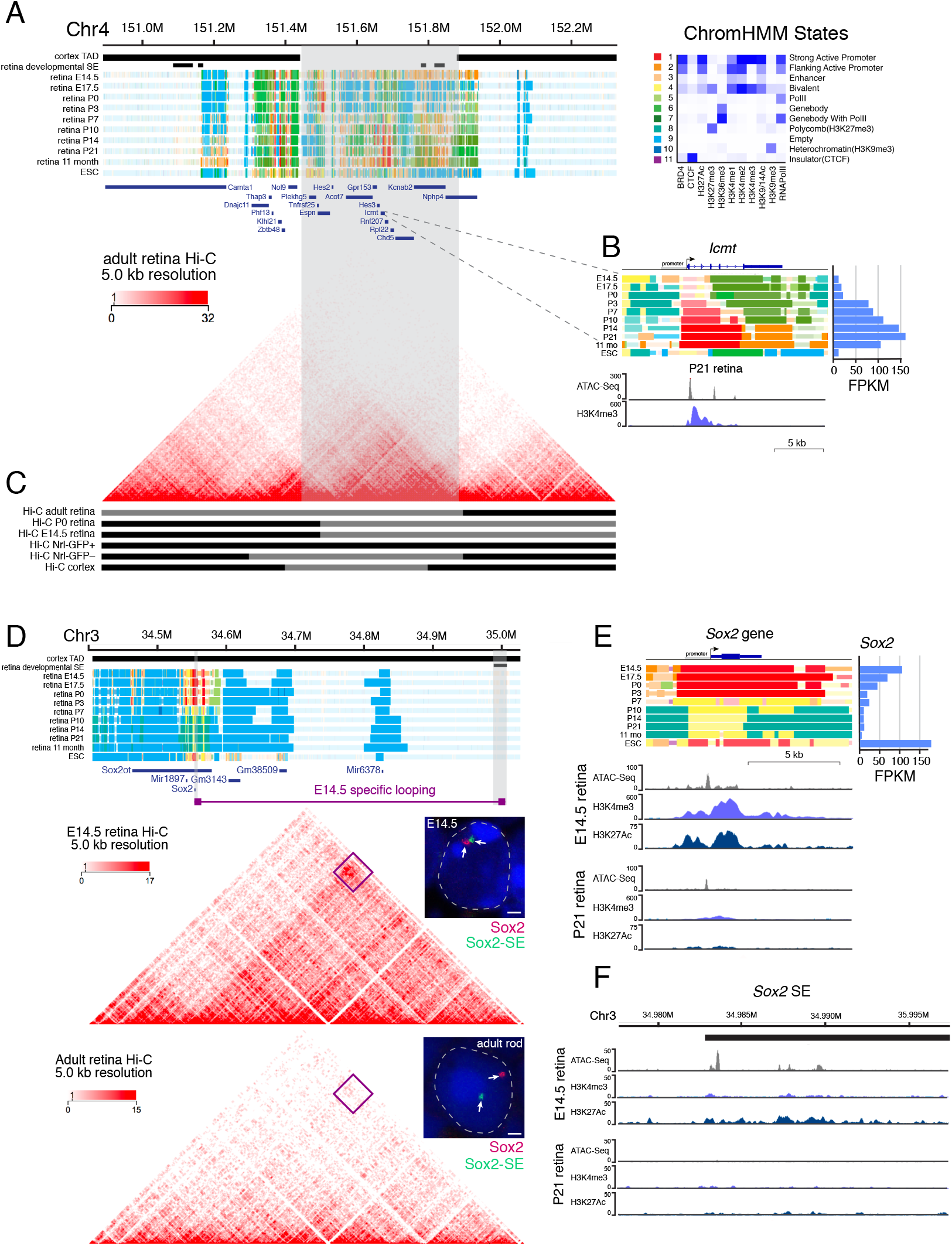
Hi-C analysis of developing retina and rod photoreceptors. **A**) ChromHMM of a retinal-specific TAD (gray-shaded region). **B**) ChromHMM and expression of the *Icmt* gene throughout retinal development with ATAC-Seq and H3K4me3 ChIP-seq track from P21 retina. Scales are indicated on the left of each track. **C**) Hi-C data of adult retina aligned with the ChromHMM data in (**A**). The gray and black bars indicate alternating topologic domains in each Hi-C dataset. **D**) ChromHMM and Hi-C data for E14.5 and adult retinae at the *Sox2* locus. The gene and associated superenhancer are highlighted (gray). An E14.5-specific predicted looping interaction from the Hi-C is indicated by the purple line and purple boxes. Two-color FISH for the *Sox2* gene (red) and superenhancer (green) is shown for a representative E14.5 retinal progenitor cell and an adult rod photoreceptor. **E**) ChromHMM, ChIP-seq, and gene expression analyses of the *Sox2* gene. **F**) ChIP-seq of the *Sox2* superenhancer. Scale bars: 1 μm. Abbreviations: Chr, chromosome; ESC, embryonic stem cells; FPKM, fragments per kilobase of million reads; SE, superenhancer

By integrating the Hi-C data with our previous CTCF ChIP-seq data, we were able to predict long-range interactions at E14.5, P0, and adult retina and in purified rod photoreceptors (Table S2). For example, a region of the genome that was more than 8 Mb away and spanned multiple TADs was predicted to be in close proximity with the rhodopsin gene *Rho* in rod photoreceptors (Fig. S1C). The long-distance interactions were confirmed by 2-color DNA FISH (Fig. S1D).

Next, we analyzed the interactions between developmentally upregulated or downregulated genes and nearby enhancers that showed similar developmental kinetics. Using the strict criteria for stable DNA-looping predictions from Hi-C data (Supplemental Information)^12^, we identified promoter–enhancer loops that during development were dynamically regulated in a manner that correlated with gene expression (Table S2). For example, there were changes in the 3D chromatin landscape of the *Sox2* promoter and a previously identified downstream super-enhancer^7^ (Fig. 1D-F). Our Hi-C data predicted a looping interaction in retinal progenitor cells expressing the gene, and that interaction was absent at later stages of development (Fig. 1D,E). Both the enhancer and promoter are inactive in the adult retina when the gene is not expressed (Fig. 1E,F). Using 2-color FISH for the *Sox2* gene and its enhancer, we discovered that the enhancer is sequestered to the facultative heterochromatin domain in rod photoreceptors, while the promoter and gene body remain in the euchromatin domain (Fig. 1D-F).

Rods contained the highest percentage of genes with developmental stage–specific promoter–enhancer looping (4.2%) and superenhancers that are activated during retinogenesis (3.9%) (Table S2)^1^. Overall, only a small proportion (1.5%) of the 4313 genes that are upregulated during retinal development showed evidence of developmental stage–specific difference in looping that correlated with the change in promoter and enhancer activity during retinogenesis (Table S2)^1^. Similar results were obtained for the 3918 genes that are downregulated during development (Table S2)^1^. None of the housekeeping genes showed any change in looping interactions during retinal development (Table S2). To determine if additional changes occurred in the promoter-enhancer interactions that did not meet our strict criteria for stable loop formation^12^, we analyzed the promoter–enhancer contacts from the Hi-C data and identified those that correlated with changes in expression (Table S2). Using this approach, we found that 25% (39/156) of rod genes, including *Rho, Crx, Aipl1, Rp1, Prom1*, and *Pde6a*, had increasing promoter–enhancer interactions that correlated with gene expression (Table S2). Similarly, 16% (38/235) of the progenitor genes, including *Pcna, Dkk3, Foxn4, Cdk1, Tuba1b*, and *Sfrp2*, had fewer promoter–enhancer contacts during retinogenesis, as the genes were silenced. Our Hi-C data identified dynamic looping interactions and more subtle differences in promoter–enhancer contacts that were associated with transcriptional changes during retinal development.

## Euchromatin and heterochromatin localization

Although nuclear size and shape can vary dramatically across cell types, the organization of heterochromatin and euchromatin is generally conserved^7^. A more detailed analysis of the heterochromatin in rods revealed a dense central core of constitutive heterochromatin with telomeres and centromeres and a more diffuse zone of facultative heterochromatin adjacent to the euchromatin domain (Fig. 2A-C)^9^. Facultative heterochromatin is thought to be cell type specific, and constitutive heterochromatin is conserved across cell types^18^.

**Figure 2.**
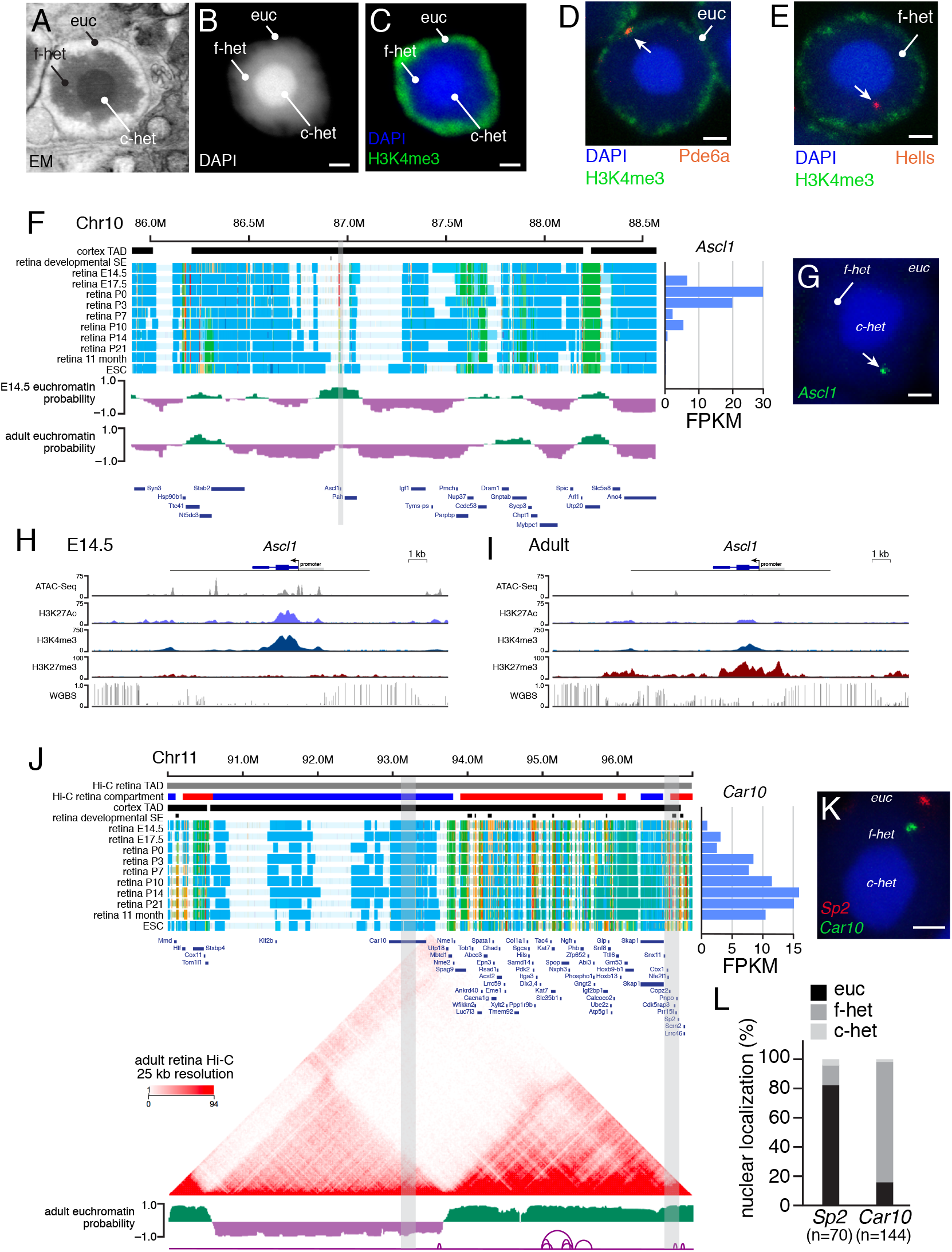
Mapping of euchromatin and heterochromatin in rod photoreceptors. **A**) Electron micrograph of a murine rod photoceptor nucleus. **B**) Confocal micrograph of a DAPI-stained rod photoreceptor nucleus. **C**) Confocal micrograph of a rod photoreceptor nucleus immunostained for H3K4me3 (green) and counterstained with DAPI (blue). **D,E**) FISH-IF for *Pde6a* and *Hells* (red) with H3K4me3 (green) and counterstained with DAPI (blue). **F**) ChromHMM of the genomic region spanning the *Ascl1* gene (gray). Euchromatin (green) and heterochromatin (purple) predictions are shown below the ChromHMM. G) FISH for *Ascl1* (green) counterstained with DAPI (blue). **H,I**) ChIP-seq and WGBS tracks for the *Ascl1* gene in E14.5 and adult retina. **J**) ChromHMM, Hi-C, and euchromatin/heterochromatin predictions for the genomic region spanning the *Car10* and *Sp2* genes (gray), which are predicted to reside in the same TAD defined from previously published cortex Hi-C. Predicted looping interactions are shown at the bottom of the panel with purple lines. The expression of the *Car10* gene is shown at the right of the ChromHMM. **K**) Two-color FISH for the *Car10* (green) and *Sp2* (red) genes. The gray bars in (J) indicate the coordinates of the FISH probes. **L**) Stacked bar plot of the quantification of *Car10* and *Sp2* localized to the rod photoreceptor domains. The numbers of nuclei scored are indicated below each bar. Scale bars: 1 μm. Abbreviations: c-het, constitutive heterochromatin; Chr, chromosome; EM, electron micrograph; ESC, embryonic stem cells; euc, euchromatin; f-het, facultative heterochromatin; FPKM, fragments per kilobase per million reads; SE, superenhancer; TAD, topological associated domain

To elucidate the subnuclear localization of developmentally regulated genes and enhancers in rod photoreceptors, we performed DNA FISH and immunofluorescence (FISH/IF) detection of H3K4me3 to mark the euchromatin domain (Fig. 2C-E). We selected constitutively expressed genes, retinal cell type-specific genes, retinal progenitor genes, silent genes, and repressed genes for our initial FISH/IF analysis (Table S3). The localization of individual loci to nuclear domains was analyzed using 3D confocal imaging and an automated image segmentation and scoring algorithm developed for retinal FISH/IF^8^ (Table S3 and Supplemental Information). Genes that are expressed in rods (e.g., *Pde6a, Rho, Crx, Nrl*, and *Nr2e3*) colocalize with H3K4me3 in the euchromatin domain (Fig. 2D and Table S3). Several retinal progenitor genes (e.g., *Ascl1, Hells, Sfrp1, NeuroD4*, and *Sox11*) were found in the facultative heterochromatin domain of rods, as were a subset of other repressed genes in our initial FISH-IF analysis (Fig. 2E and Table S3).

To extend these observations to the entire genome, we developed a machine learning–based algorithm to predict euchromatin/heterochromatin localization (Supplemental Information). The model was trained using the FISH/IF data for 103 loci (Table S3) and genomic features extracted from the chromHMM data^1^. Initially, the modeling was constrained to TAD boundaries from the mouse cortex^11^. This provided a robust prediction of TAD-based euchromatin/heterochromatin status with an accuracy of 89% (10-fold cross validation, Supplemental Information). Inclusion of additional genomic (LINE/SINE repeats), epigenomic (WGBS, ATAC-seq) or transcriptome (RNA-seq) data did not significantly improve performance (89% accuracy).

Most TADs are either euchromatin or heterochromatin, but we identified some that were a mixture of both. Therefore, we removed the TAD-boundary constraint in our modeling, which achieved a similar performance (87% accuracy) (Table S3). To independently validate the euchromatin/heterochromatin predictive model, we performed FISH for an additional 161 loci and achieved 89% accuracy using the regional level model learned from the entire dataset. In total, 22 of 264 loci were discordant between the FISH-IF and the computational modeling. Nine of the discordant loci contained large genes (>100 kb) expressed at low levels such that the promoter had chromHMM features consistent with euchromatin, but the gene body had features consistent with heterochromatin due to its large size and empty chromHMM state (Table S3). In addition, 14 of 22 discordant loci were at the boundary between euchromatin and heterochromatin domains (Table S3). The predictive model has been extended to E14.5 retinae and added to the cloud-based viewer, where researchers can compare the 1D (histone and DNA modifications), 2D (looping interactions), and 3D (nuclear subdomain localization) nuclear organization for any genomic region. Using the model, we found that virtually all rod and housekeeping genes are predicted to be localized to the euchromatin domain. However, 14% (32/246) of the retinal progenitor genes and 53% (307/692) of the non-rod cell type specific genes (e.g. *Grik2, Gria3, Ctn4, Epha5, Cav1*) are predicted to be localized to the heterochromatin domain of rods. Similarly, 21% (88/410) of cancer genes (e.g. *Myc, Myb, Foxp1, Kit, Cdkn2a, Cdk6*) and 13% (37/277) of stress response genes (e.g. *Hspa13, Dnajc5b, Nell, Eno1b*) are predicted to be localized to the heterochromatin domain of rods.

The aforementioned example of the *Sox2* enhancer transitions during development from euchromatin to heterochromatin (Fig. 1), and several other retinal progenitor genes, such as *Hells* and *Ascl1*, transition from euchromatin in retinal progenitor cells to heterochromatin in rods (Fig. 2F-I). We also identified genes, such as *Rp1*, that transition from heterochromatin to euchromatin during retinal development (Fig. S2). Similarly, there is evidence that cell type-specific localization of genes to heterochromatin is important for differentiation (Tables S3,S4). For example, *Car10* is expressed from the euchromatin domain in bipolar neurons^19^ but is sequestered in the facultative heterochromatin domain of rods (Tables S3,S4 and Fig. 2J-L). This locus is another example of retinal-specific chromatin-domain organization. The *Car10* and *Sp2* genes are predicted to reside in a single TAD from the murine cortex Hi-C data but not the retinal Hi-C data; euchromatin/heterochromatin prediction and FISH show they are in different domains (Fig. 2J-L). In total, 6.2% (246/3918) of genes that are downregulated during retinal development were sequestered to heterochromatin in the adult retina, and 9.3% (4/43) of G2/M-phase cell cycle genes and 4.6% (11/235) of retinal progenitor cell genes showed the same pattern (Tables S3,S4). Among the genes that are upregulated during retinal development, 7.6% (331/4313) were predicted to be in heterochromatin in E14.5 retinal progenitor cells; 7.3% (14/190) of rod genes showed a similar pattern. The most striking result was that 15.4% (82/533) of the genes that are upregulated during retinal differentiation and have corresponding changes in chromHMM started out in the heterochromatin domain in E14.5 retinal progenitor cells and transitioned to euchromatin in adult retina (Tables S3, S4 and Fig. S2).

## Inversion of nuclear organization

Our integrated dataset showed that some genomic loci undergo developmental transitions in their histone and DNA modifications, DNA looping, and subnuclear localization. However, many regions are more stable throughout development. To determine if the physical association of heterochromatin with the nuclear lamina is essential for proper gene regulation in the retina, we performed RNA-seq analysis on Lamin B receptor (Lbr)-mutant mouse retinae. In the developing mouse retina, the Lbr protein helps tether heterochromatin to the nuclear periphery in newly postmitotic cells^20^. A 2-bp insertion in the *Lbr* gene (*Lbr^ic-J^*) has been previously described: homozygous-mutant mice have severe developmental defects and die around P8-P12^21^. Heterozygous mutations in the human *LBR* gene are associated with the Pelger-Huët anomaly, which is characterized by defects in nuclear morphology and chromatin distribution^22^. Occasionally, some *Lbr*^*ic-J*/*ic-J*^ mice survive to adulthood, but the retinae have not been analyzed in those animals^20^.

We harvested retinae from wild-type, *Lbr^ic-J^*-heterozygous and -knockout mice at P12 and maintained those retinal explants in culture for 2 weeks to complete retinal differentiation. In addition, we obtained 1 litter of mice that survived to P21 and included those retinae in our analysis. We identified a significant increase in the amount of heterochromatin that was not tethered to the nuclear lamina in the inner nuclear layer (INL) of *Lbr*^*ic-J*/*ic-J*^ mice (Fig. 3A,B). The areas of the nucleus and individual domains of heterochromatin that remained tethered to the nuclear lamina in the INL cells of the *Lbr*^*ic-J*/*ic-J*^ mice did not significantly differ from those in their wild-type littermates (Fig. 3C,D). However, the area of individual untethered heterochromatin domains was significantly increased in the *Lbr*^*ic-J*/*ic-J*^ mice (Fig. 3E; p=0.042). As expected, the organization of euchromatin and heterochromatin in rods was unaffected because those cells do not express Lbr^20,23^ (Fig. 3A). RNA-seq analysis of P21 *Lbr*^*ic-J*/*ic-J*^, *Lbr*^+/*ic-J*^, and *Lb*^+/+^ littermates demonstrated that very few genes were deregulated in the absence of *Lbr* (Table S5). Together, these data suggest that euchromatin and heterochromatin domains are determined as part of the cell type-specific differentiation programs in the retina, and changes in attachment to the nuclear lamina has little impact on gene expression.

**Figure 3.**
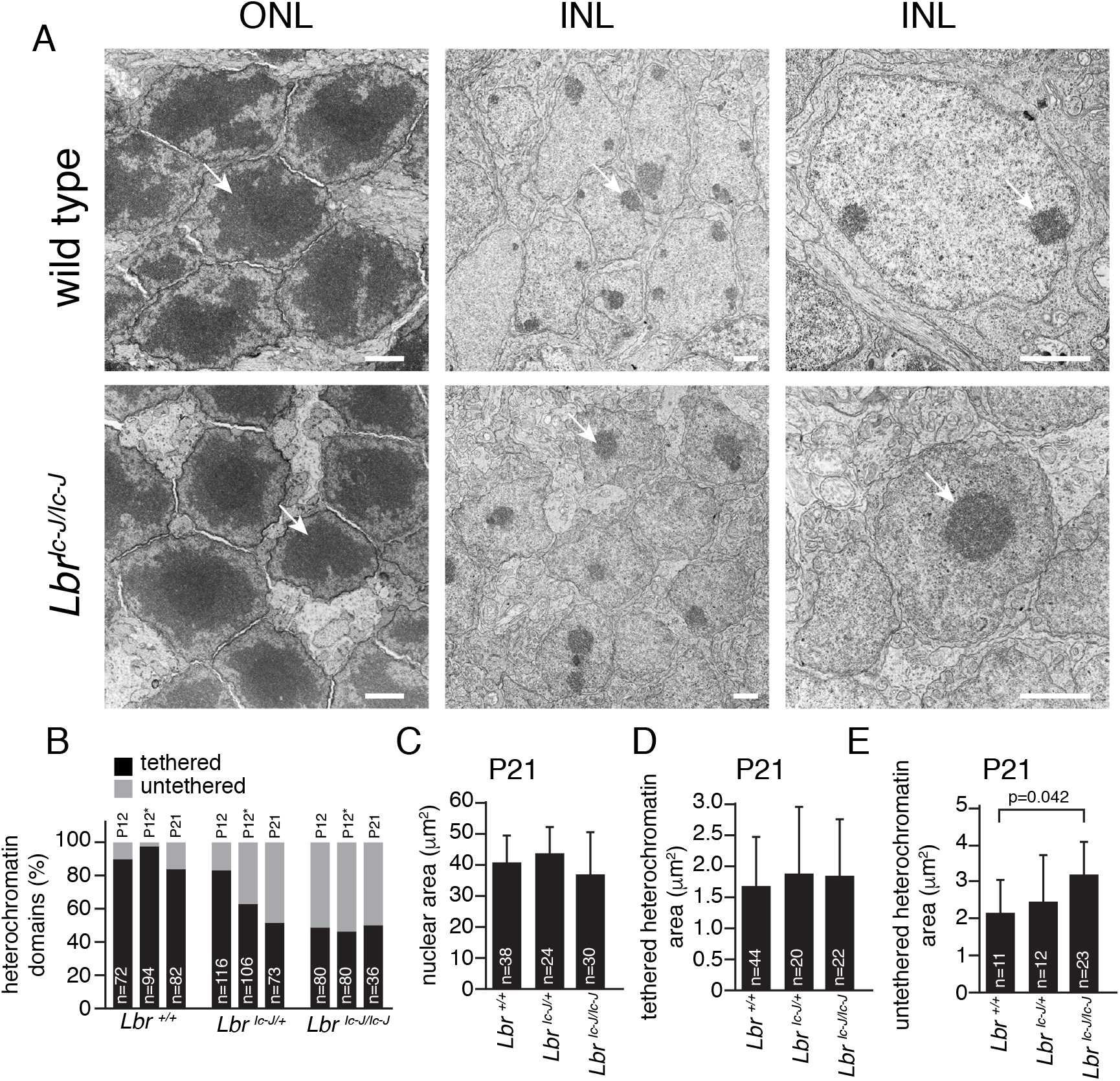
Untethered heterochromatin in *Lbr*-deficient retinae. **A**) Electron micrographs of wild-type and *Lbr*^*ic-J*/*ic-J*^ mouse retinae. Arrows indicate the heterochromatin domain in rods found in the outer nuclear layer (ONL) and neurons and glia found in the inner nuclear layer (INL). **B**) Stacked bar plot of the percentage of heterochromatin domains that are tethered or untethered to the nuclear lamina. The number of domains scored is indicated on each bar. The (*) indicates scoring on samples that were harvested at P12 and maintained in culture for 9 days. **C-E**) Bar plots of nuclear area (C), area of tethered heterochromatin domains (D), and untethered heterochromatin domains (E) from electron micrographs for *Lbr*^+/+^, *Lbr*^+/*ic-J*^, and *Lbr*^*ic-J*/*ic-J*^ INL cells. The means and standard deviations are plotted, and the numbers of nuclei scored are indicated on each bar. Scale bars: 1 μm.

## Cell type–and developmental stage-specific core regulatory circuit super-enhancer

To validate our integrated genomic dataset and visualization tools in vivo, we selected a gene with a complex expression pattern during retinal development. The paired-type homeobox gene *Vsx2* is expressed in retinal progenitor cells throughout development, and mutations in *Vsx2* lead to microphthalmia in humans and mice^24–27^. In addition, Vsx2 is expressed in mature bipolar neurons and at lower levels in Müller glia^28,29^. It is not known how this precise temporal and spatial pattern of expression is achieved during retinal development. Previous studies have shown that a bacterial artificial chromosome spanning the *Vsx2* gene and 55 kb of upstream sequence is sufficient to confer retinal progenitor, bipolar, and Müller glial cell expression of a reporter gene in transgenic mice^28,29^. Other studies have identified a proximal promoter element that contributes to retinal progenitor cell expression and an upstream element sufficient for bipolar neuron expression^29,30^ (Fig. 4A). However, none of those elements were shown to be required in vivo. We discovered a CRC-SE that was developmental stage specific and consistent with bipolar cell expression of Vsx2 (Fig. 4A). Indeed, the region within the CRC-SE that is most conserved across species contains a Vsx2 consensus-binding site (Fig. 4B-F).

**Figure 4.**
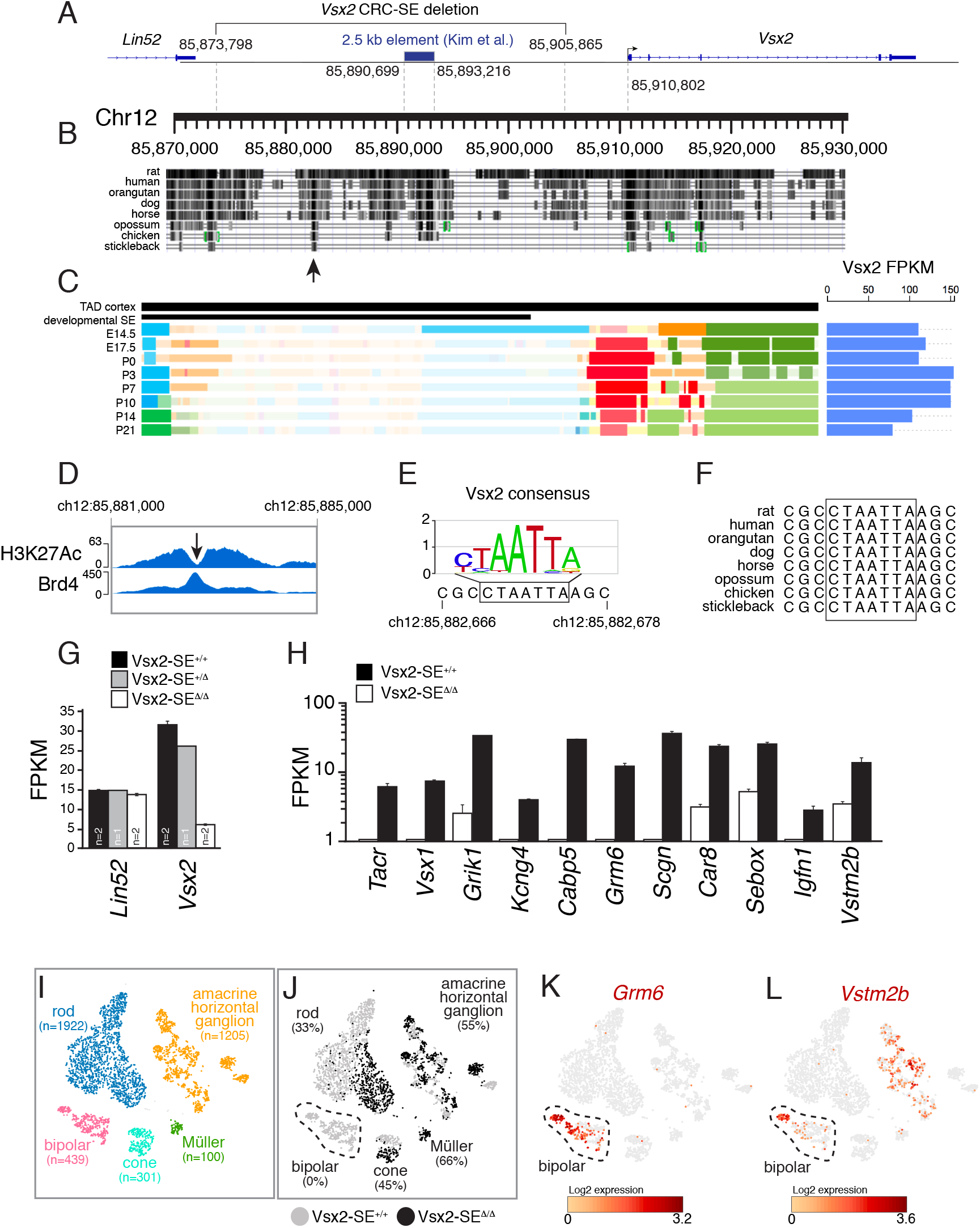
Identification of a bipolar neuron–specific CRC-SE. **A**) Genomic map of the murine *Vsx2* locus showing a 2.5-kb region previously identified in heterologous reporter assays as containing candidate genes for the bipolar neuron–specific regulatory element. The 32-kb putative *Vsx2* CRC-SE is indicated with genomic coordinates from mm9. **B**) Evolutionary conservation across the *Vsx2* locus with an arrow indicating a region conserved down to fish (stickleback). **C**) ChromHMM and RNA-seq of the *Vsx2* gene during development showing the developmental stage-specific superenhancer (black bar). **D**) ChIP-seq for H3K27Ac and Brd4 for the evolutionarily conserved region indicated by the arrow in (**B**). **E,F**) The arrow in (**D**) indicates a trough in H3K27Ac that has a Vsx2 consensus-binding site. **G**) Bar plot of gene expression from RNA-seq analysis of *Lin52* and *Vsx2* for the *Vsx2-SE*^+/+^, *Vsx2-SE*^+/*Δ*^, and *Vsx2-SE^Δ/Δ^* retinae. Each bar is the mean and standard deviation for the number of animals indicated. **H**) Bar plot of gene expression from RNA-seq analysis of 11 representative bipolar neuron–specific genes that span subtypes of bipolar neurons. Each bar is the mean and standard deviation of biological replicates. (**I**) T-distributed stochastic neighbor embedding (tSNE) plot of singlecell RNA sequencing for 1 adult *Vsx2-SE*^+/+^ and a *Vsx2-SE^Δ/Δ^* littermate. Cell types were assigned based on known gene expression profiles from previous studies. **J**) Distribution of cells to the different cell type-specific clusters based on genotype of the retina that was analyzed. The percentage of each cell type cluster that was from the *Vsx2-SE^Δ/Δ^* retina is indicated. **K,L**) Heatmap of gene expression overlaid on the tSNE plot from (**I**) for a bipolar neuron–specific gene that is completely absent in the *Vsx2-SE^Δ/Δ^* retinae (*Grm6*) and another that is only partially downregulated *(Vstm2b).* The bipolar cell cluster in the tSNE is circled. Abbreviations: Chr, chromosome; FPKM, fragments per kilobase per million reads; SE, superenhancer; TAD, topological associated domain

To determine if the conserved region is required for bipolar neuron-specific expression, we deleted the CRC-SE in mice by using CRISPR-Cas9. In total, 3 independent sublines were made that could be distinguished by their deletion junctions (Supplemental Information). The individual sublines were intracrossed to produce litters with wild-type (*Vsx2-SE*^+/+^), heterozygous (*Vsx2-SE*^+/*Δ*^), or homozygous (*Vsx2-SE*^*Δ*/*Δ*^) deletion of the *Vsx2* CRC-SE. RNA-seq confirmed the significant downregulation of *Vsx2*, as well as other bipolar neuron-specific genes^31^ (Fig. 4G,H and Table S6). To extend our bulk RNA-seq analysis, we performed singlecell transcriptome analysis of an adult *Vsx2-SE*^+/+^ and *Vsx2-SE^ΔΔ^* retinae (Fig. 4I,J). Within the resolution of our Drop-Seq experiment, all bipolar cells were absent in the *Vsx2-SE^Δ/Δ^* retinae. Genes that are expressed solely in bipolar neurons such as *Grm6* (rod bipolar cells) were absent in the RNA-seq analysis of *Vsx2-SE^Δ/Δ^* retinae (Fig. 4H,K and Table S6). However, genes expressed in other cell typrs in addition to bipolar neurons such as *Vstm2b* had residual expression in the RNA-seq analysis of *Vsx2-SE^Δ/Δ^* retinae (Fig. 4H,L and Table S6).

There was no evidence of microphthalmia in the *Vsx2-SE^Δ/Δ^* mice, which was consistent with data showing that the CRC-SE was active only when bipolar cells were generated (Fig. 5A,B). We measured visual acuity using the OptoMotry system and found that the *Vsx2-SE^Δ/Δ^* mice lacked any optomotor response under bright light (cone) conditions (Fig. 5C). We then immunostained retinal vibratome sections from each of the 3 genotypes using antibodies specific for proteins found in rods (rhodopsin), cones (cone arrestin), bipolar neurons (PKCα, G_o_α, Vsx2), Müller glia (glutamine synthetase), reactive Müller glia (GFAP), horizontal neurons (calbindin), and amacrine cells (Pax6, calretinin, TH). Bipolar cells were absent and the outer plexiform layer (OPL) was disrupted with enlarged cone pedicles (Figs. 5D,E and S3). This phenotype resembled retinae from a recent study that disrupted Wnt5a/b signaling between photoreceptors and bipolar neurons at the OPL and is consistent with the loss of bipolar neurons^32^. There was no evidence of Müller glial cell reactive gliosis from the immunostaining or the RNA-seq analysis (data not shown). Transmission electron microscopy confirmed the absence of bipolar cell bodies, and 3D electron microscopy using the Helios focused ion beam scanning electron microscope (FIB/SEM) was used to reconstruct rod and cone terminals in the *Vsx2-SE*^+/+^ and *Vsx2-SE^Δ/Δ^* mice. In total, 509 10-nm sections were collected and analyzed from the *Vsx2-SE*^+/+^ OPL and 490 10-nm sections were collected and analyzed from the *Vsx2-SE^Δ/Δ^* OPL. Across 30 rod spherules and 6 cone pedicles, we found no evidence of bipolar dendrites in the *Vsx2-SE^ΔΔ^* retina (Fig. 5F-I). Together, these data indicate that the deletion of a developmental stage– and cell type-specific CRC-SE led to the complete absence of bipolar cells in the murine retina.

**Figure 5.**
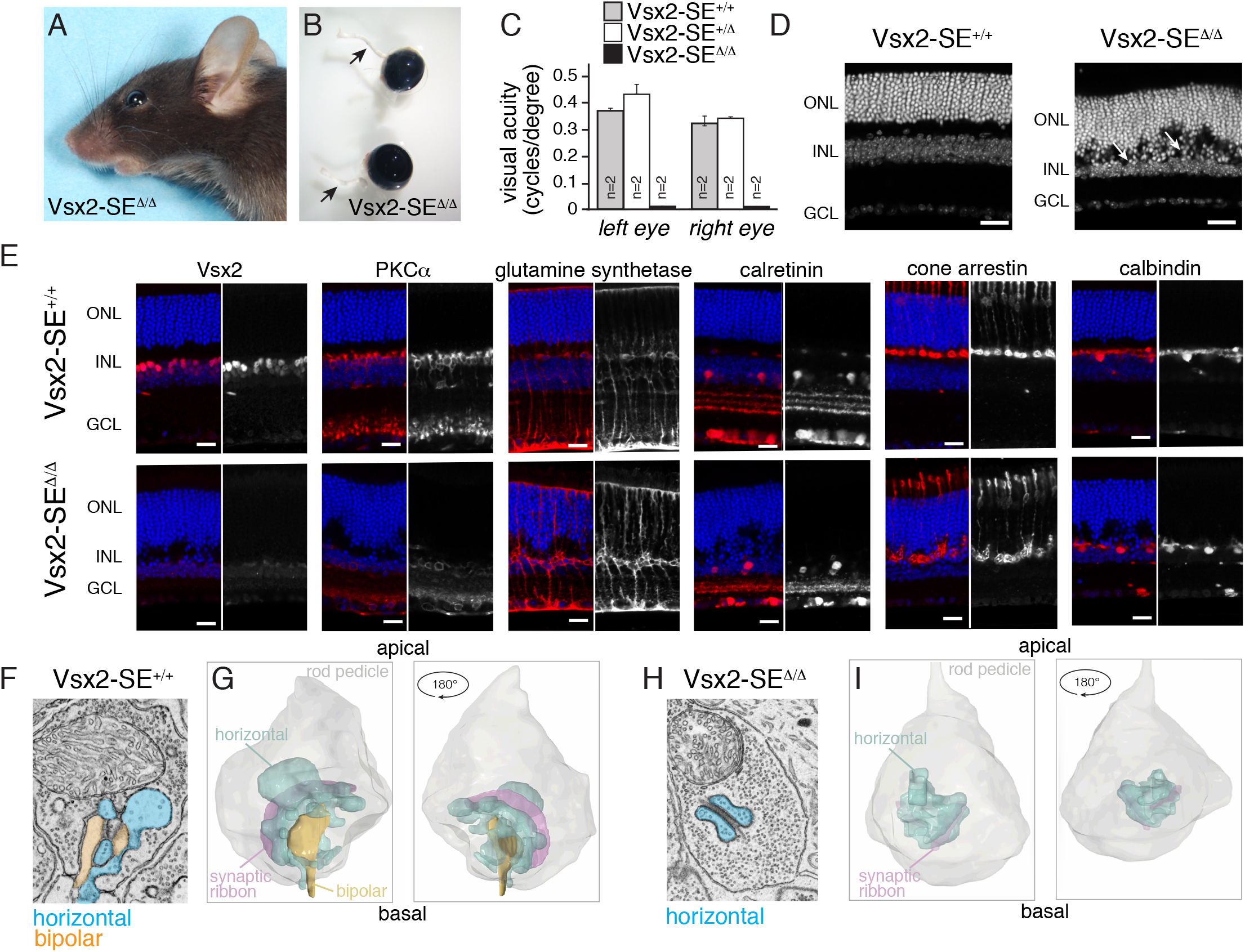
Mice lacking the *Vsx2* CRC-SE are blind due to the loss of bipolar neurons. **A,B**) Photograph of a *Vsx2-SE^Δ/Δ^* mouse head (**A**) and eyes (**B**). Arrows indicate the optic nerves. **C**) Bar plot of visual acuity measured by the optomotor response for 8-week-old *Vsx2-SE*^+/+^, *Vsx2-SE*^+/*Δ*^, and *Vsx2-SE^Δ/Δ^* mice. Data represent the means and standard deviations for 2 mice for each genotype measured on 4 successive days. **D**) Confocal micrographs of DAPI-stained adult retinae. Arrows indicate disruption in the outer plexiform layer. **E**) Confocal micrographs of adult retinae immunostained using antibodies for specific retinal cell types. Side-by-side images with the immunostaining (black and white) and DAPI overlay (color) are shown. **F,G**) Representative electron micrographs (**F**) and 3D tracing (**G**) of rod spherules from Vsx2-SE+^/^+and *Vsx2-SE^Δ/Δ^* mice (**H,I**). The horizontal cell processes can be identified by the presence of synaptic vesicles (blue); bipolar dendrites (orange) lack synaptic vesicles. Scale bars: 25 μm. Abbreviations: GCL, ganglion cell layer INL, inner nuclear layer; ONL, outer nuclear layer.

## DISCUSSION

Here we built upon our previous 1D epigenetic profiling of the murine retina to include 2D looping interactions predicted by Hi-C analysis, 3D chromatin landscape profiled by FISH/IF and computational modeling in developing and adult retinal cells. Our data revealed longdistance interactions that can span multiple TADs and complex multigene interactions, which is consistent with transcriptional hubs for cell type-specific gene expression. This highlights the value of integrating Hi-C data with ChIP-seq, DNA methylation, and transcriptome data to identify developmentally regulated, tissue-specific chromatin domains.

We also showed that important developmental, cancer, and stress-response genes are localized to the facultative heterochromatin in rods, which have a unique inverted nuclear structure in mice. Our integrated analysis was used to model 3D localization in rods providing the first 1D, 2D and 3D nucleome model for any neuronal cell type. Subnuclear localization of genes to facultative heterochromatin is a tightly regulated and cell type-specific process. However, untethering heterochromatin from the nuclear lamina in the retinal neurons and glia that have a conventional nuclear architecture did not significantly affect gene expression. This suggests that the cell type-specific organization of the genome into heterochromatin and euchromatin domains is separate from the physical distribution of those domains within the nucleus. Thus, the nuclear size, shape, and distribution of heterochromatin and euchromatin may be more relevant for cell and tissue biology than for gene expression per se. Our integrated online visualization tool was useful for identifying a developmental stage– and cell type-specific CRC-SE upstream of *Vsx2*, and we showed that this element was required for expression of *Vsx2* in bipolar neurons in vivo. With similar profiling of other cell types, it should be feasible to generate the same type of 1D/2D/3D map of virtually any normal or diseased cell type.

## MATERIALS AND METHODS

### Hi-C methods

In situ Hi-C protocol was performed using retinal cells, with minor modification to the original protocol^1^. Two to five million cells were crosslinked with 1% formaldehyde for 10 min at room temperature, then digested with 125 units of MboI overnight followed by labeling with biotinylated nucleotides and proximity ligation. After reverse crosslinking, ligated DNA was purified and sheared to 300-500 bp using a Covaris LE220. The DNA fragments containing ligation junctions were pulled down with streptavidin beads followed by Illumina compatible library construction and paired end sequencing.

### Confocal image analysis of FISH probes

Images were taken with Zeiss LSM 700 confocal microscope using 63X lens. To map FISH probe nuclear position, we employed machine-learning methods ^10^ to segment different nuclear region types (core: blue channel, DAPI-ring: blue channel and Green ring: green channel, FISH spots: red channel) in each 2D plane of the z-stacks. The machine-learning classifier uses 85 feature images for each plane that are created based on texture, gray-scale intensity and neighborhood information. We then use the 3DROIManager ImageJ plug-in ^11^ that calculates common surface area between the segmented regions in the 3D segmented image. We use the values of common surface area to identify FISH spots inside each region.

## ACKNOWLEDGEMENTS

We thank Angie McArthur for editing the manuscript. This work was supported, in part, by Cancer Center Support (CA21765) from the NCI, grants to M.A.D from the NIH (EY014867 and EY018599 and CA168875) and ALSAC. M.A.D. was also supported by a grant from Alex’s Lemonade Stand Foundation for Childhood Cancer, the Tully Family Foundation, and the Peterson Foundation. The majority of this research was supported by the Howard Hughes Medical Institute.

---

**Figure S1.**
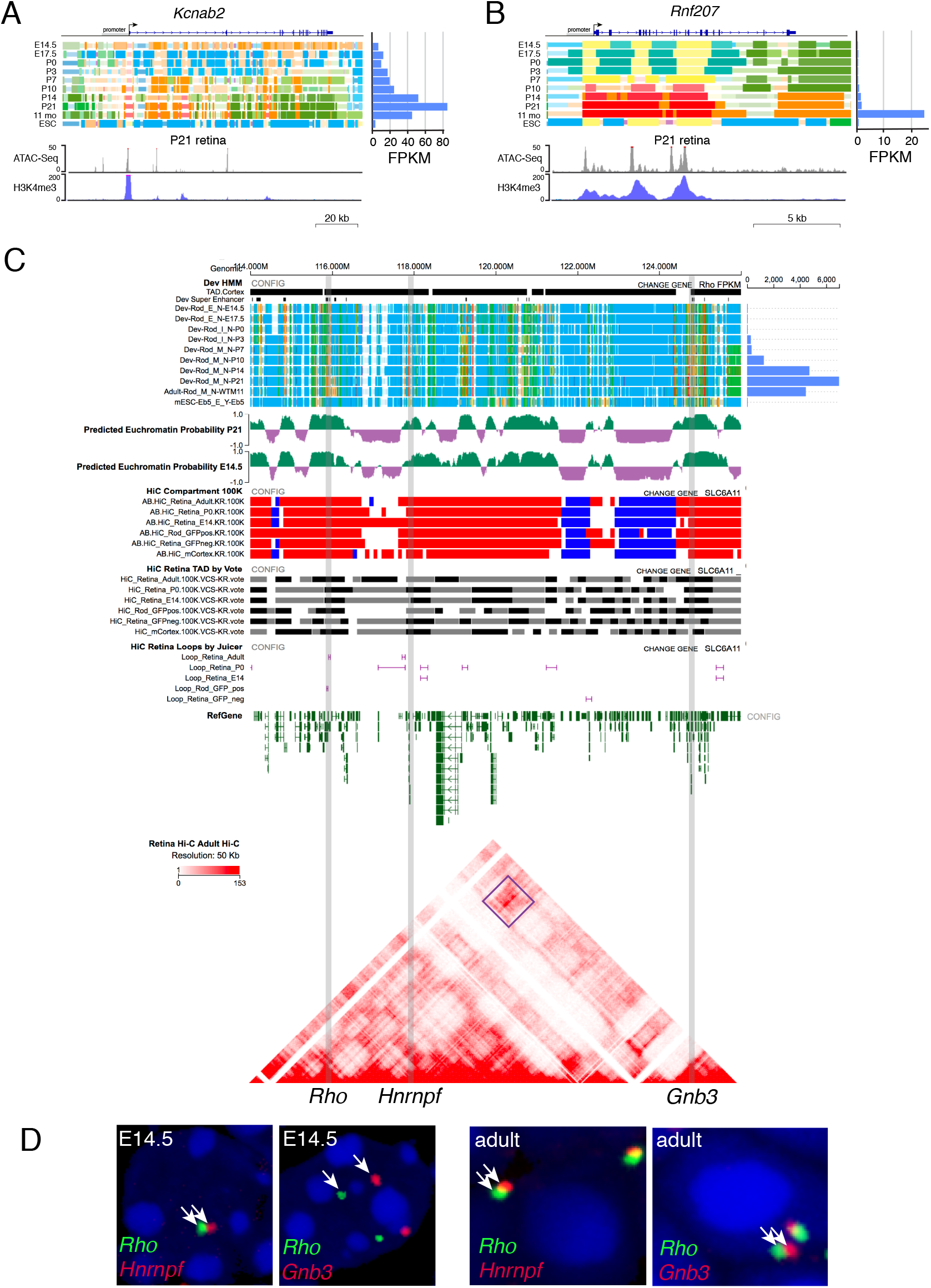
Long-range interactions with the *Rho* gene locus. **A,B**) ChromHMM of the *Kcnab2* and Rnf207 genes with gene expression (RNA-seq) showing increased expression during development. ATAC-seq and H3K4me3 ChIP-seq tracks are shown below the gene. **C**) ChromHMM, euchromatin/heterochromatin prediction, Hi-C compartmentalization, gene looping and Hi-C data for a 12 Mb region including the *Rho, Hnrnpf* and *Gnb3* genes. **D**) Micrographs of 2 color FISH for Rho (green) and Hnrnpf (red) or Gnb3 (red) in E14.5 retinal progenitor cells or adult rod photoreceptors.

**Figure S2.**
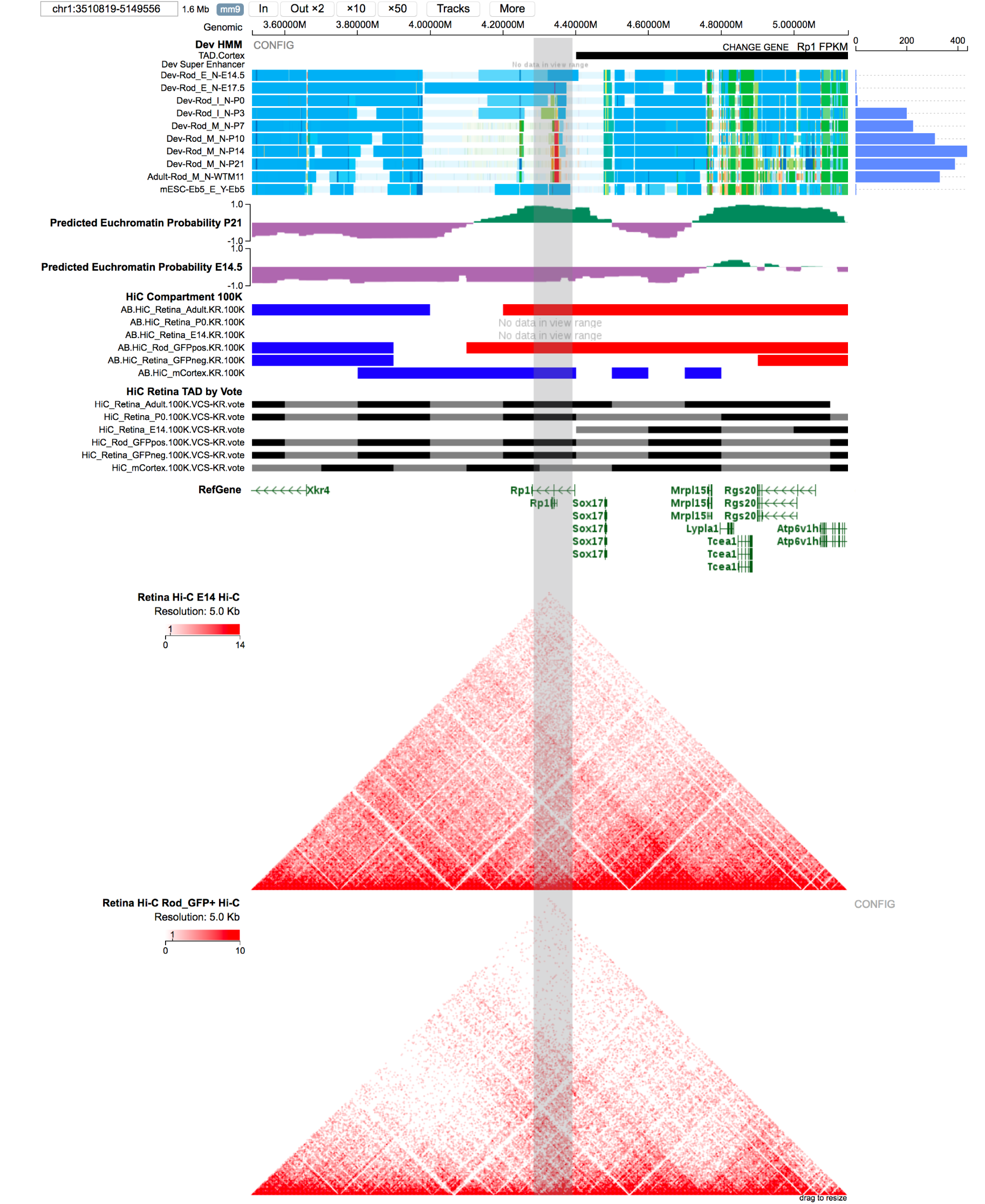
Euchromatin/heterochromatin dynamics during development. ChromHMM, RNA-seq, euchromatin/heterochromatin prediction and Hi-C data for the *Rp1* gene at E14.5 and adult retina. In E14.5 retinae, the *Rp1* gene is predicted to localize to heterochromatin while in the adult retina it is in the euchromatin.

**Figure S3.**
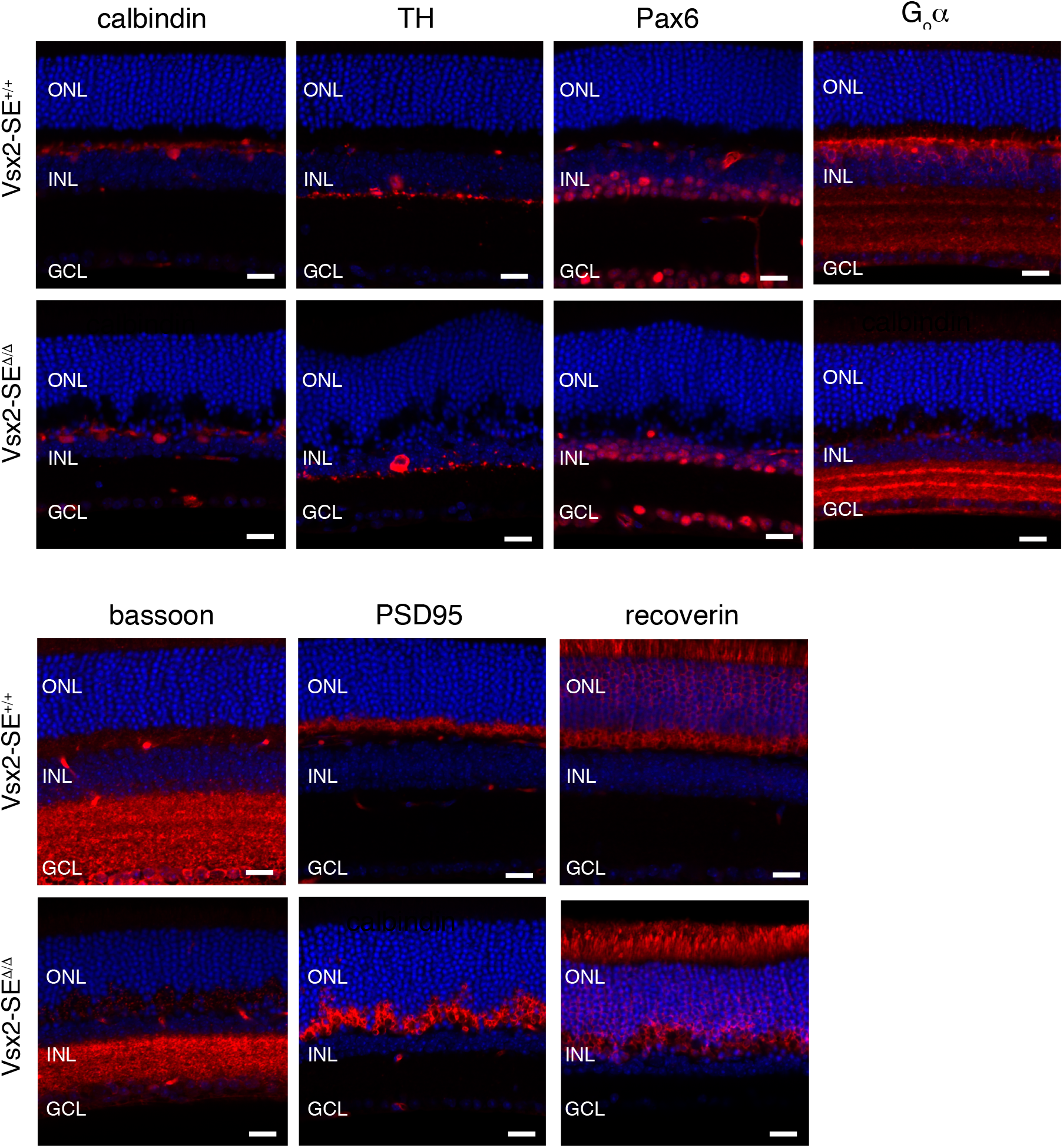
Immunostaining of *Vsx2-SE^Δ/Λ^* retinae. Additional immunofluorescence for horizontal cells (calbindin), dopaminergic amacrine cells (TH), amacrine cells (Pax6), bipolar cells (G_o_α), synapses (bassoon and PSD95) and photoreceptors (recoverin). The sections were counterstained with DAPI (blue). Scale bars: 25 μm.

## Hi-C data analysis

All HiC data have been processed by the Juicer pipeline (version 1.5)^2^ against mouse genome mm9. One of our adult mouse retinal samples was sequenced on the Illumina GAIIx and all other samples were sequenced on the Illumina Novaseq. We confirmed there were no sequencing artifacts before merging the files from the two platorms using HiC-spector (version 201706028 from github)^3^. Mouse cortex raw data were downloaded from GEO: GSE35156. All 5 samples had more than 4.9 billion contacts and 3 samples had >7.5 billion contacts providing a resolution of 1 kb. Stable loops were identified using HiCCUPS from Juicer at 5kb and 10kb resolution.

The highest quality (MAPQ >30), 370~640 loops were called from each sample and those that were less than 3 Mb were included in the visualization online.

Computational analysis of TAD boundaries is dependent on the computational pipeline^4^. We chose Armatus because it had the best overlap of TAD boundaries and CTCF binding sites in our dataset. The TADs called by Armatus using both normalized method(KR:Knight-Ruiz, VC_SQRT:square root of vanilla coverage) had a similar trend of increased TAD number over development. 100K resolution achieved the best balance between TAD number and the enrichment of CTCF sites at TAD boundaries. We also noticed some large regions enriched for Hi-C contacts that were not called as TADs either by KR or VC_SQRT, so we developed a “vote” mechanism to merge the TADs called by KR to VC_SQRT only if the TADs called by KR did not overlap with TADs called by VC_SQRT.

## Hi-C data analysis

All HiC data have been processed by the Juicer pipeline (version 1.5)^1^ against mouse genome mm9. One of our adult mouse retinal samples was sequenced on the Illumina GAIIx and all other samples were sequenced on the Illumina Novaseq. We confirmed there were no sequencing artifacts before merging the files from the two platorms using HiC-spector (version 201706028 from github)^2^. Mouse cortex raw data were downloaded from GEO: GSE35156. All 5 samples had more than 4.9 billion contacts and 3 samples had >7.5 billion contacts providing a resolution of 1 kb. Stable loops were identified using HiCCUPS from Juicer at 5kb and 10kb resolution. The highest quality (MAPQ >30), 370~640 loops were called from each sample and those that were less than 3 Mb were included in the visualization online.

Computational analysis of TAD boundaries is dependent on the computational pipeline^3^. We chose Armatus because it had the best overlap of TAD boundaries and CTCF binding sites in our dataset. The TADs called by Armatus using both normalized method(KR:Knight-Ruiz, VC_SQRT:square root of vanilla coverage) had a similar trend of increased TAD number over development. 100K resolution achieved the best balance between TAD number and the enrichment of CTCF sites at TAD boundaries. We also noticed some large regions enriched for Hi-C contacts that were not called as TADs either by KR or VC_SQRT, so we developed a “vote” mechanism to merge the TADs called by KR to VC_SQRT only if the TADs called by KR did not overlap with TADs called by VC_SQRT.

## An epigenetic-based classifier for euchromatin/heterochromatin domain prediction

We integrated global epigenetic signatures of multiple histone marks and transcription factors of retina cells at various developmental stages and explored the feasibility of predicting the euchromatin/heterochromatin domain status from epigenetic and transcriptomic features. Although the experimental euchromatin mark (H3K4me3) was included in the epigenetic modeling, our chromHMM model estimated that only 1.80% (range: 1.47-2.15%) of the genome have strong H3K4me3 signal (States 1 and 2), which is substantially lower from the experimentally estimated fraction of euchromatin domains. Therefore, H3K4me3 alone is not sufficient to classify euchromatin and heterochromatin on a genome-wide scale.

Previous studies have found that TADs overlap with linear chromatin domains defined by histone modifications and TAD boundaries are somewhat conserved across cell types. As a first step, we evaluated the performance of a predictive model using features extracted from chromHMM modeling to determine the euchromatin/heterochromatin status of predefined TADs (based on mouse cortex). The analysis of 103 genomic regions that were analyzed by FISH suggested that ChromHMM based features provided accurate and robust prediction of TAD-based euchromatin/heterochromatin status (accuracy: 0.89 +/- 0.09 in 10-fold cross validation). Inclusion of additional genomic/epigenomic/transcriptomic features did not improve performance (0.89 +/- 0.09). The euchromatin/heterochromatin prediction improved to 0.95 when we removed the TAD boundary constraint and this is consistent with sub-TAD regions of euchromatin and heterochromatin. On an independent validation set of 161 genes/regions we achieved an accurace of 0.89.

Based on the accuracy of the original model, we next generated a model based on all 264 (103+161) genes/regions collected in P21 retinal cells, which was used to derive the genomewide probability of being euchromatin using ChromHMM features in a sliding-window of 200kb with a step size of 20kb. We further applied the p21 model to epigenetic data collected on E14.5 retina cells to evaluate whether the classifier is robust in data collected independently. Although the selected regions were biased towards euchromatin domains in both training (68%) and validation (70%) sets, the genomwide prediction revealed that majority of genome found were in heterochromatin regions for both E14.5 (62.9%) and P21 (65.4%) cells. P21 cells packaged more genomic regions to heterochromatin domains. The predicted euchromatin/heterochromatin topology is consistent with the high-resolution HiC data, providing independent validation of the modeling data. Further result suggested that while most TADs were dominated by one chromatin type (heterochromatin or euchromatin), occasionally these two domains co-existed in a single TAD. Moreover, although most TADs are relatively stable among different developmental stages, a subset of them displayed dynamic boundaries.

## ChromHMM features

For a defined region (TAD or sub-TAD region) in the genome, 11 features were derived, representing the fraction of the region covered by each of the 11 chromatin states.

## Other TAD level features analyzed

1. Distance from nearest TAD boundary
2. TAD size
3. Epigenetic state of TAD
4. Distance from nearest expressed gene (FPKM > 10)
5. Distance from nearest enhancer
6. LINE repeat density at gene + promoter
7. SINE repeat density at gene + promoter
8. LINE repeat density in region
9. SINE repeat density in region
10. ATAC-Seq peak density at gene + promoter
11. ATAC-Seq peak density in region
12. Expression level of targeted gene
13. DNA methylation of targeted gene (promoter and whole gene)

We generated an ensemble classifier combining three component models: Linear SVM ^5^, PCA Logistic Regression L2 ^6,7^, and Random Forest ^8^. For each component, a model was trained to predict the probability of a given region in euchromatin. The prediction from the ensemble classifier is the average of the individual probabilities of each component. The algorithms were implemented in Sci-Kit Learn ^9^.

Using the 103 training genes/regions, we evaluated three feature sets: 1) All Features, 2) Features based only on TAD_HMM [1-11] and 3) features without TAD_HMM [1-11]. We performed 10-Fold Cross Validation and report the results of the three feature sets.

| Feature Set | CV Accuracy (mean +/- stdev) |
| --- | --- |
| ALL | 0.885 +/- 0.092 |
| ChromHMM features | 0.885 +/- 0.092 |
| ALL without chromHMM features | 0.827 +/- 0.098 |

Using the 103 training genes/regions, we trained two ensemble classifiers, one based on TAD-level ChromHMM features (for TAD-based classification) and the other based on region-level ChromHMM features (for sub-TAD region based classification). The performance of the classifiers was evaluated on a set of independent validation set of 161 genes/regions, not seen in the classifier building step.

We generated a final classifier using all 264 genes/regions (combined from training and validation sets) in P21 retina cells and predicted the probability of being in euchromatin for genomwide regions inboth P21 and E14.5 retina cells using a sliding window of 200kb with a step size of 20kb.

## 3DEM fixation and processing

The samples for 3DEM were fixed in 2% paraformaldehyde and 2.5% glutaraldehyde in 0.1M cacodylate buffer overnight and rinsed in same buffer. The tissue was stained using a modified heavy metal staining method (Ellisman et al.) and processed through a graded series of alcohol, propylene oxide and infiltrated in propylene oxide /epon gradients. The tissue was then infiltrated overnight in 100% Epon 812 resin and polymerized for 48 hours in a 70°C oven.

## Preparation steps for focused ion beam scanning electron microscopy imaging

The sample block is mounted on an aluminum pin stub with a conductive silver epoxy, and sputter-coated with a thin layer (~60nm) of iridium to electrically ground the sample reducing charging. The block face is scanned to locate the ROI. The block is trimmed to the ROI using an ultramicrotome with a diamond knife. Next a relief is milled into the block face using a suitable ion beam current, and a protective cap is deposited using the carbon gas injection system. Fiducials are created to aid the computer vision pattern placement algorithm.

## Focused ion beam scanning electron microscopy

The samples are imaged and processed for 3D data collection on a Helios G3 system. Imaging is normally at 2kV, 100pA, 5×5×5nm, ICD. Regions of interest are imaged utilizing Auto Slice and View 4 software package to automate serial sectioning and data collection processes.

## Scoring Lbr-deficient heterochromatin

Images of retinae were taken on the Tecnai T20 and heterochromatin aggregate attachment was scored for each nucleus as tethered or untethered based on contact with the nuclear lamina. Nuclei and chromatin aggregates in the inner nuclear layer were traced by hand, and their areas quantified by ROI measurement in Fiji. Nuclei bearing classic features of horizontal cells or Müller glia were excluded.

## Scoring of Vsx2-CRC-SE knockout

Individual images from a sequential, block-face of the outer retina from wild type (a total 509 images) and knockout samples (a total of 490 images) were analyzed. For analysis of spherules, three areas, each containing cross sections of a minimum of 10 spherules, provided a total of 30 spherules each for scoring wild type and knockout samples. For each of the three areas, the thirty adjacent images containing the full set of cross sections of the ten selected spherules were then examined in order to determine the presence or absence of synaptic vesicles within all invaginating processes. Because of the relatively large size of pedicles, the entire series of sequential, block-face images was used to select six representative examples of pedicles from each wild type and knockout sample. All adjacent images containing the full set of cross sections of pedicles from the six wild type and six knockout samples were analyzed for presence or absence of synaptic vesicles within all invaginating processes.

## Combining immunofluorescence protein staining with DNA FISH

Freshly cut unfixed cryosections were fixed in 1% PFA in PBS for 5 minutes, followed by an additional fixation in 1% PFA and 0.05% NP40. Fixed samples were stored in 70% ethanol at -20^0^C until needed. Following fixation slides were subjected to immunostaining by first blocking in 1% BSA and 2X SSC for 10 minutes followed by protein detection using various antibodies at appropriate dilutions in the same blocking solution. Detections were carried out at RT for 45 minutes followed by washing in PBS for 5 minutes. Appropriate secondary antibodies were applied using the same procedure as for primary antibodies. Slides were then fixed in 4% PFA and 0.5% Tween 20 and 0.5% NP40 for 10 minutes, followed by treatment in 0.2N HCl containing 0.5% Triton X-100 for 10 minutes. Slides were then denatured in 70% formamide and 2X SSC at 80^0^C for 10 minutes. Following denaturation, the slides were dehydrated in a graded ethanol series consisting of 70%, 80%, and 100% ethanol for 2 minutes each. Denatured probes were then applied to the slides under 22 × 22 mm coverslips at 37^0^C overnight in a solution containing 50% formamide, 2X SSC, and 10% dextran. Slides were then washed in 50% formamide and 2X SSC at 37^0^C for 5 minutes followed by mounting in Vectashield containing DAPI.

## Confocal image modification

Brightness and contrast were modified for images presented in the figures for the FISH/IF studies. Raw original data are available for all datasets and probes.

## Crispr-CAS9 mediated deletion of Vsx2-CRC-SE

A total of 65 pups were generated from 5 separate days of injections. On each day, ten 3-4 week old C57BL/6J female mice from Jackson Labs were superovulated with 5 units of gonadotrophin from pregnant mare’s serum (PMSG from ProSpec) and 48 hours later, with 5 units of human chorionic gonadotrophin (hCG from Sigma). After overnight mating with C57BL/5J males, the females were euthanized and oocytes were harvested from the ampullae. The protective cumulus cells were removed using hyaluronidase, and the oocytes were washed and graded for fertilization by observing the presence of two pronuclei. A mixture provided by the Center for Advanced Genome Engineering of 100 ng/ul Cas9 mRNA and 25-50 ng/ul each of two guide RNAs, ZZ30.mVsx2.g7 (5’ GCAGGCCATGTGCTCGTCGA 3’) and ZZ31.Vxs2.g16 (5’ CAGGGTGCAGGCTGACAACG 3’), was injected into the cytoplasm of the oocytes. gRNAs were designed to have at least 3bp of mismatch between the target site and any other site in the mouse genome. gRNAs were tested prior to embryo injection for activity in mouse N2A cells using targeted next generation sequencing as previously described^12^. They were then returned to culture media (M16 from Millipore or A-KSOM from Millipore) and later the same day transferred to day 0.5 pseudopregnant fosters (7-10 week old CD-1 females from Charles River Laboratories mated to vasectomized CD-1 males). Pups were born after 19 days gestation and genotyped using PCR and Illumina Mi-Seq. Positive animals were weaned at day 21 and at 6 weeks of age, they were back-crossed to C57BL/6J mice. The primers used for genotyping are shown below with Mi-Seq adaptors (red):

> Forward:
>
>
> > 5’-TCGTCGGCAGCGTCAGATGTGTATAAGAGACAGGCTCTGACCTTCCTGGAAGCCCCGC-3’
>
> Reverse:
>
>
> > 5’-GTCTCGTGGGCTCGGAGATGTGTATAAGAGACAGCTCAGGAGGTTACAAGGAGGTGTAG-3’

The primers span 32 kb and will not give a PCR product for the wild type allele but will give a 300 bp product in the deleted allele. We also generated a set of primers internal to the deletion to serve as a wild type primer set to distinguish heterozygous and homozygous deleted mice. Those internal wild type primers will produce a PCR product of 149 bp and are:

> Forward:
>
>
> > 5’-CATAACTGGCTGTATTCTGTGTGACTC-3’
>
> Reverse:
>
>
> > 5’-CTTACATCCTTTGACCCTGGCTATG-3’

The deletion junctions for the 3 sublines are:

> Vsx2-3-3: 5’-TCTAC/CCTGCACCCTGAAATCA-3’

> Vsx2-59-12: 5’-TCTACCCT/CACCCTGAAATCA-3’

> Vsx2-23-3: 5’-TCTACCCT/GCACCCTGAAATCA-3’

## RNA-Seq

RNA was extracted from each iPS cell lines by using RNeasy Plus Mini kit (Qiagen, 74134) or Direct-zol kit (Zymo Research, R2050). Libraries were prepared from ~500 ng total RNA with the TruSeq Stranded Total RNA Library Prep Kit according to the manufacturer’s directions (Illumina). Paired-end 100-cycle sequencing was performed on HiSeq 2000 or HiSeq 2500 sequencers according to the manufacturer’s directions (Illumina.)

## 10x Genomics single cell sequencing

To dissociate the retinal tissue, 40 U papain was added to 400 μl papain buffer (1mM L-cysteine with 0.5mM EDTA in PBS -/-) and incubated at 37°C for 30 min to activate the enzyme. 1 adult retina was added to the tube and incubated for 20-30 minutes with gently agitation every 10 minutes until the tissue dissociated. 40 μl of DNase and 40 μl of soybean trypsin inhibitor were added per tube. The cells were gently triturated with a P1000 barrier pipette tip 8 times and incubated for 5 min in a 37 °C water bath. The cells suspension was transferred to a 50 ml conical bottom tube through a 40-μm mesh cell strainer and the strainer was PBS^-/-^ to bring the total volume to 3 ml. 5 ml of BSA cushion (4% BSA in retinal explant medium) was placed in a 15 ml conical bottom tube and the 3 ml cell suspension was gently overlaid onto the BSA cushion. The tube was centrifuged at 500xg for 10 minutes at 4 °C to pellet the cells. The supernatant was aspirated and the cells were resuspended in 400 μl of explant culture medium.

The 10x Genomics Chromium controller with the version 2 library and gel bead kit was used along with the i7 multiplex index kit (120262). Single Cell A Chip Kit (120236) and Single Cell 3’ Library & Gel Bead kit version 2 (120237) were used in this protocol. Sequencing was done on an Illumina HiSeq4000 and the alignment was done on cell ranger (v2.1.1) using mm10 genome reference. Data were visualized using Loupe Cell Browser (v2.0.0).

## Optomotry

The OptoMotry system from CerebralMechanics was used to measure the optomotor response of Vsx2-SE^Δ/Δ^ mice. Briefly, a rotating cylinder covered with a vertical sine wave grating was calculated and drawn in virtual three-dimensional (3-D) space on four computer monitors facing to form a square. Vsx2 mice standing unrestrained on a platform in the center of the square tracked the grating with reflexive head and neck movements. The spatial frequency of the grating was clamped at the viewing position by repeatedly recentering the cylinder on the head. Acuity was quantified by increasing the spatial frequency of the grating until an optomotor response could not be elicited. Contrast sensitivity was measured at spatial frequencies between 0.1 and 0.45 cyc/deg.

In total, 2 Vsx2-SE^Δ/Δ^, 3 Vsx2-SE^+/Δ^ and 2 Vsx2-SE^+/+^ were analyzed along with 2 C57Bl/6J mice. The tester was blinded to genotype until after testing was complete. Mice were acclimated to the device for 1 week and then tested on 4 successive days the following week.

## Immunostaining

For immunostaining, retinae from 9.4 week old littermates were isolated and fixed in 4% PFA overnight at 4°C. Retinae were washed 3x with PBS and then embedded in 4% LMP agarose in PBS for vibratome sectioning at 50 μm. They were incubated in block solution for 1 hr at room temperature and then in primary antibody in block solution overnight at 4°C.

The antibodies were:

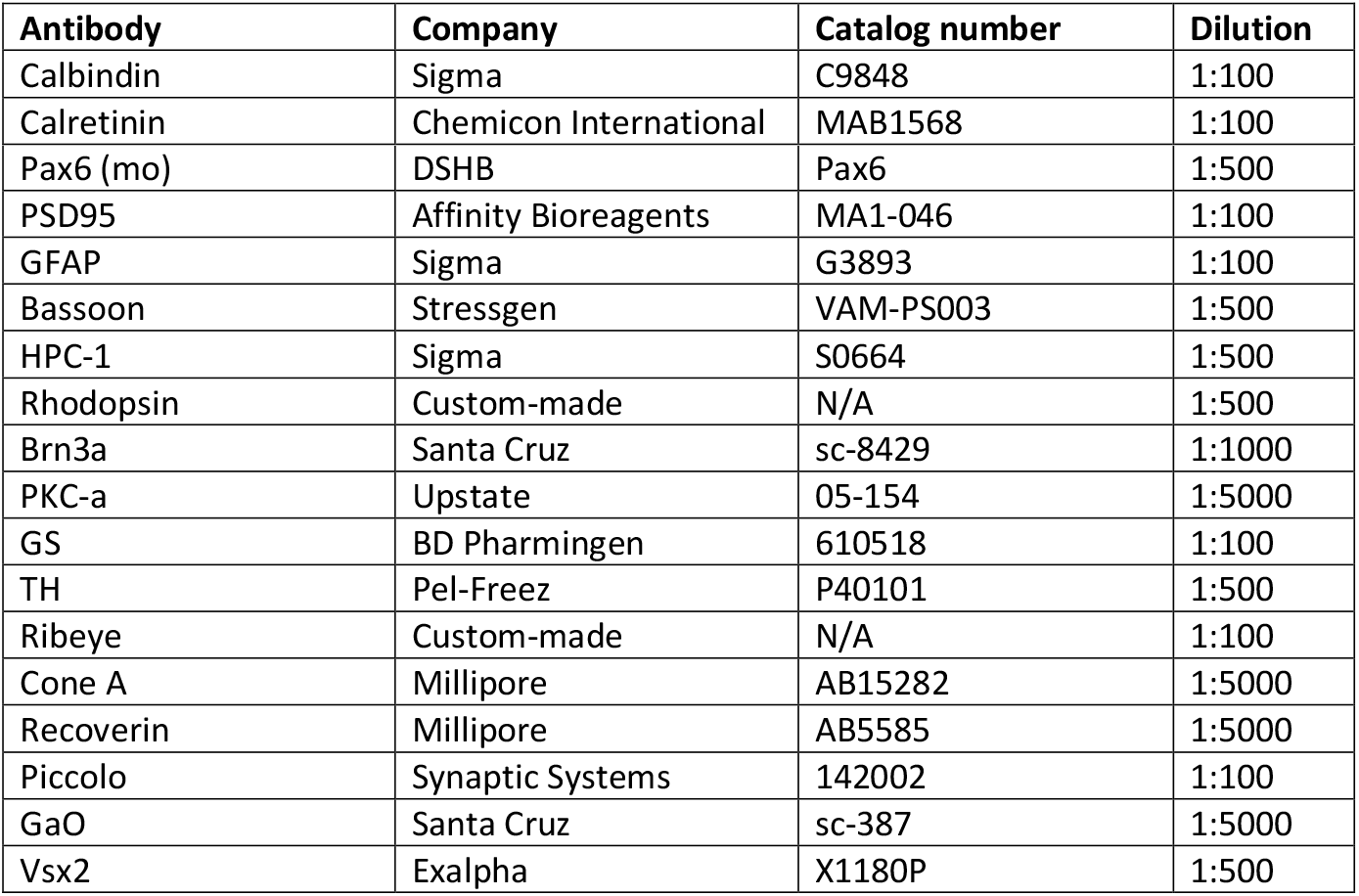

| Antibody | Company | Catalog number | Dilution |
| --- | --- | --- | --- |
| Calbindin | Sigma | C9848 | 1:100 |
| Calretinin | Chemicon International | MAB1568 | 1:100 |
| Pax6 (mo) | DSHB | Pax6 | 1:500 |
| PSD95 | Affinity Bioreagents | MA1-046 | 1:100 |
| GFAP | Sigma | G3893 | 1:100 |
| Bassoon | Stressgen | VAM-PS003 | 1:500 |
| HPC-1 | Sigma | S0664 | 1:500 |
| Rhodopsin | Custom-made | N/A | 1:500 |
| Brn3a | Santa Cruz | sc-8429 | 1:1000 |
| PKC-a | Upstate | 05-154 | 1:5000 |
| GS | BD Pharmingen | 610518 | 1:100 |
| TH | Pel-Freez | P40101 | 1:500 |
| Ribeye | Custom-made | N/A | 1:100 |
| Cone A | Millipore | AB15282 | 1:5000 |
| Recoverin | Millipore | AB5585 | 1:5000 |
| Piccolo | Synaptic Systems | 142002 | 1:100 |
| GaO | Santa Cruz | sc-387 | 1:5000 |
| Vsx2 | Exalpha | X1180P | 1:500 |

Vibratome sections were washed twice in PBS and then incubated in secondary antibody for 1 hour at room temperature. All secondary antibodies were incubated a dilution of 1:500 in the appropriate block solution. We used donkey anti-mouse (Vector Labs BA-2000), goat anti-rabbit (Vector Labs BA-1000) and rabbit anti-sheep (Vector Labs BA-6000). After secondary antibody, they were washed twice with PBS and and incubated with ABC reagent (Vector Laboratories, Cat. # PK6100) for 30 minutes. We then used tyramide Cy3 (PerkinElmer, Cat. # FP1046) for 10 minutes at room temperature and washed 2x in PBS followed by DAPI at 1:1000 in PBS. Slices were mounted in Prolong gold reagent and imaged on a Zeiss LSM700 confocal microscope.

